# Democratising “Microscopi”: a 3D printed automated XYZT fluorescence imaging system for teaching, outreach and fieldwork

**DOI:** 10.1101/2020.05.21.108894

**Authors:** Matthew Wincott, Andrew Jefferson, Ian M. Dobbie, Martin J. Booth, Ilan Davis, Richard M. Parton

## Abstract

Commercial fluorescence microscope stands and fully automated XYZt fluorescence imaging systems are generally beyond the limited budgets available for teaching and outreach. We have addressed this problem by developing “Microscopi”, an accessible, affordable, DIY automated imaging system that is built from 3D printed and commodity off-the-shelf hardware, including electro-mechanical, computer and optical components. Our design features automated sample navigation and image capture with a simple web-based graphical user interface, accessible with a tablet or other mobile device. The light path can easily be switched between different imaging modalities. The open source Python-based control software allows the hardware to be driven as an integrated imaging system. Furthermore, the microscope is fully customisable, which also enhances its value as a learning tool. Here, we describe the basic design and demonstrate imaging performance for a range of easily sourced specimens.

**Highlights:**

- Portable, low cost, self-build from 3D printed and commodity components
- Multimodal imaging: bright field, dark field, pseudo-phase and fluorescence
- Automated XYZt imaging from a tablet or smartphone via a simple GUI
- Wide ranging applications in teaching, outreach and fieldwork
- Open source hardware and software design, allowing user modification

## INTRODUCTION

Microscopes are essential tools in many fields of science. However, in recent times the basic microscope stand has evolved into an automated imaging system, with a corresponding escalation in complexity and cost. In order to inspire and educate the next generation of budding scientists and to make modern microscopy widely available, wherever it is needed, it is essential to facilitate access to these advances. Particularly important is to address the need for software controlled integrated hardware and high-quality fluorescence digital image capture, not typically found in portable or classroom microscopes. Recently there has been a great drive towards democratised technology, whereby advanced technology becomes increasingly available for low-budget projects. This movement has provided an impetus for the design of low-cost microscopes and the release of designs in open-source formats. The rise of 3D printing epitomises this trend and opens possibilities for the democratisation of the production process. Moreover, the falling cost of digital cameras as used in smartphones and the recent availability of compact, cheap computing power, exemplified by the Raspberry Pi (https://www.raspberrypi.org/about/) and Arduino (https://www.arduino.cc) eco-systems, has changed the landscape for delivering low cost microscopes.

To date, a number of low-cost functional microscopes have been introduced that fulfil different purposes. The implementations of such systems vary significantly in size, cost, robustness, optical quality, imaging modalities, and degree of automation. Many of these low-cost microscopes are designed to connect to smartphones, which are now almost ubiquitous. Such microscopes are frequently intended for use in global health and therefore tailored towards their applicability for field diagnostics. Early implementations employed standard objectives at the expense of an increase in bulk [Breslauer 2009]. Ultra-low cost and portable designs are typically based on a simple Leeuwenhoek design and incorporate ball lenses as the principal optical element [Bogoch 2013, Orth 2018]. Other designs incorporating two lenses have been introduced to better match a conventional microscope [Hergemoller 2017]. Furthermore, alternative imaging modalities have been demonstrated in these smartphone-based systems, including fluorescence [Zhua 2013, Wei 2013, Cybulski 2014, Switz 2014] and polarised light microscopy [Casey 2015]. Alternative approaches include that of the Foldscope [Cybulski 2014], which is formed from folded paper, costs a few dollars and again incorporates a ball lens. These solutions offer excellent portability and ultra-low cost but suffer due to aberrations inherent at the edge of the field of view due to the ball lenses employed. Although elegantly simple, the above solutions offer limited functionality and are far removed from the modern automated imaging system found in a typical research lab.

A second class of low-cost microscopes aims to replace commercial systems. These low-cost systems often make use of 3D printing in the design and can include varying degrees of functionality and mechanical or electronic control. An excellent robust 3D printed design, which could replace more expensive basic teaching microscopes, is that of Tristan-Landin *et al*., (2019), offering both bright field and fluorescence imaging on a CCD camera. However, this system lacks automation and image quality is limited by the choice of objective lens. A major challenge in recreating an automated imaging system at a low cost is in the implementation of an XYZ translation of the sample relative to the optical path which is both robust and able to cope with a variety of specimens. An example of a robust, low cost automated microscope incorporating XYZ translation is the “Open Flexure Microscope” developed by Collins *et al*., (2020). The highly compact configuration of this system, whilst excellent for portability, rapid manufacture, and use in the field, restricts convenient exploration and manipulation of the component parts. Diederich *et at*., (2020) have developed an adaptable imaging system (UC2) based on 3D printed cubes with embedded magnets. This provides a cheap, robust framework to fulfil simple fluorescence imaging requirements in the research lab, for example in tissue culture rooms, and great potential for flexibility and development. With its appealing “click together” assembly, the UC2 imaging system has the potential to provide a real alternative to expensive research microscopes in a range of situations but may be limited in its application to a wider range of imaging subjects.

Here we present Microscopi, an affordable, portable, automated DIY imaging system developed for use in interdisciplinary teaching and outreach purposes at universities and schools. The instrument is tailored to this role with easily assembled hardware, a clear layout, and a straightforward graphical user interface underpinned by well documented customisable Python-based code. Here, we document the build process and demonstrate the utility of Microscopi on a range of samples and imaging modalities. Our overarching goal is to cultivate a community of users by freely distributing detailed assembly instructions, control software and practical guides. Microscopi has further potential applications where lab facilities are unavailable due to location or limited resources.

## RESULTS and DISCUSSION

### Designing and Building a Simple 3D printed Motorised XYZ Digital Imaging System

Our main goal in developing Microscopi was to design a widely accessible, customisable, automated imaging system (Figure 1; Supplemental Figure S1) to inspire the next generation of scientists. The use of DIN, non-infinity corrected objective lenses, low-cost electronics and predominantly 3D printed parts reduces the construction cost to less than £300 per basic unit, or £400 with fluorescence capability. This cost represents good value for money, given that Microscopi incorporates automated sample navigation, digital image acquisition and processing, all controlled through a simple customisable software-interface. Sample navigation comprises a motorised Z-objective lifter (focus) and a motorised XY flexure stage delivering automated image tiling. Furthermore, we have incorporated optics supporting contrast enhancing bright field modes and fluorescence imaging of various samples. Our open design facilitates building and supports customisation of both hardware and software. The small footprint and robust design, incorporating an optional battery powerpack, allows the unit to be fully portable for use in outreach events and fieldwork. Different objective lenses may be used to adjust magnification and resolution and multiple imaging modalities allow Microscopi to cope with a variety of specimens.

**Figure 1:**
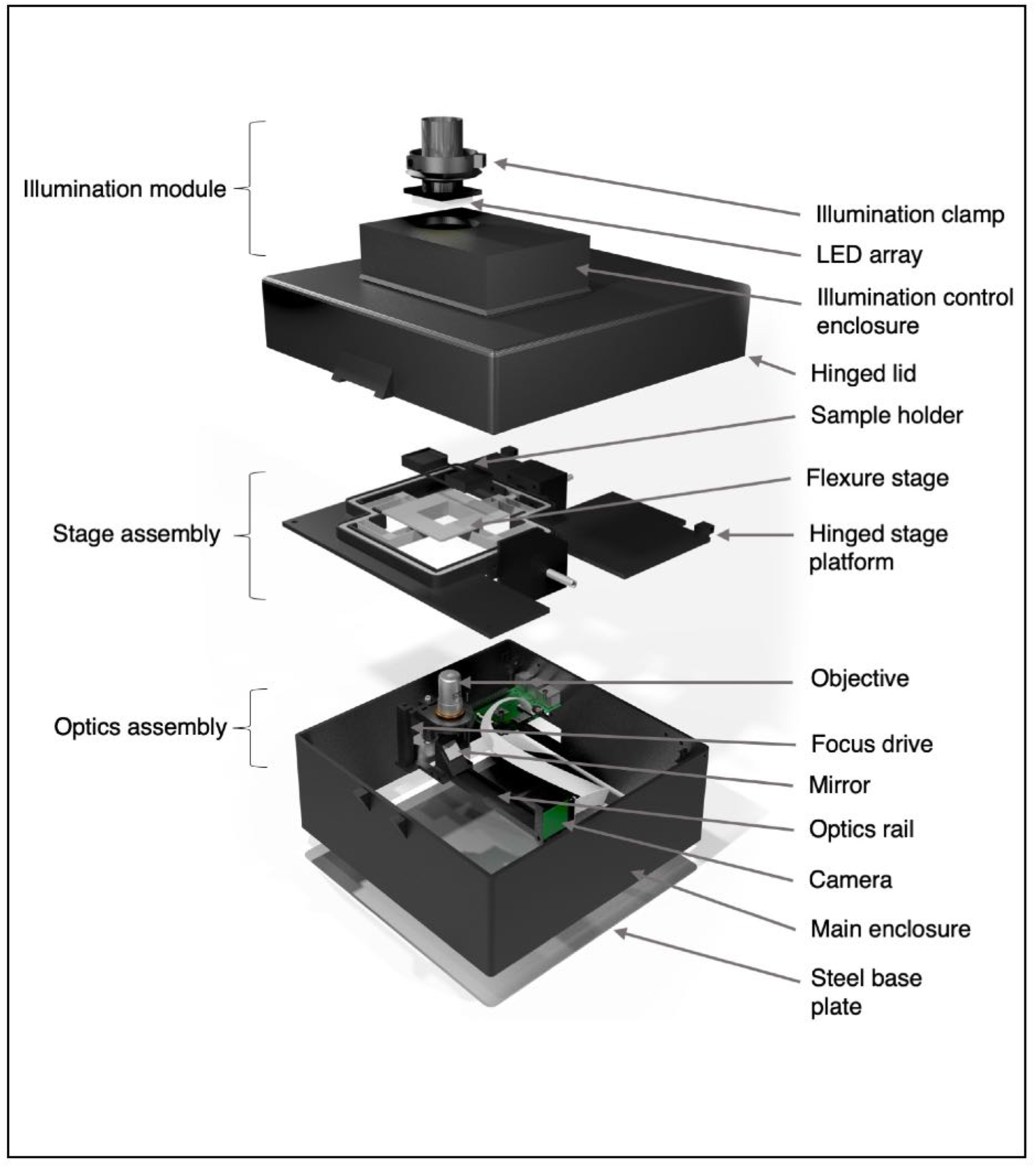
The Microscopi imaging system. Schematic diagram of the Microscopi unit demonstrating the principal assemblies. The illumination module can be switched out for modules supporting different imaging modalities. The stage assembly incorporates the motorised XY sample stage. The lid and stage assembly are hinged to allow easy access to the optics assembly and electronics. All the component parts and the assembly of the instrument are detailed in the Supplementary Methods.

### Design Specifications

- Dimensions (235 mm × 235 mm base; 250 mm height)
- Weight 1.5 kg
- 3D printed components made from PLA (Polylactic Acid)
- XY flexure stage travel 3 mm in X and Y
- Z-objective lifter range 4 mm
- Onboard processor running Python 3 (Raspberry Pi 4 Model B)
- CMOS camera, Sony IMX219 8-megapixel sensor (Pi-camera V2)
- Power: 5 V, < 4.0 A
- Portable battery run time approximately 10 h
- Optical path 160-DIN lens compatible, 45 mm parfocal distance

Assembly and operating instructions for Micoscopi, including a full listing of the commercially obtained and 3D printed parts and details of the operating software may be found online at https://doi.org/10.5281/zenodo.3701602. The software code can be found on GitHub at https://github.com/micronoxford/microscopi. A brief summary of materials is provided here in the Table of Resources. Components of Microscopi to be 3D printed were designed in a freely available CAD software package (https://www.openscad.org). Designs were exported as STL format files (“stereolithography” or “Standard Triangle Language”. Components were printed on various 3D printers, including a Lulzbot Taz 5 and an Ultimaker S5, one of the latest FDM (fused deposition modelling) printers. After printing, parts were finished using a safety razor blade, needle files set and sandpaper to remove minor printing imperfections and to smooth interlocking surfaces.

### Multiple Optical Paths for Contrast Enhancement of Diverse Samples

A key requirement in microscopy is to generate contrast from transparent material by translating features of the sample into intensity or colour differences that can be more easily visualised. To achieve this, Microscopi incorporates multiple possible optical paths for contrast generation with a variety of different kinds of samples.

Microscopi is an inverted microscope, i.e. the objective lens is situated below the sample. An inverted configuration facilitates the loading of sample material, particularly with life-science applications such as organisms in pond water. The objective lenses used are finite conjugate DIN lenses [Simon &Comastri, 2005] which focus directly onto an image sensor, providing a conceptually simple low-cost design. To assist in first setting up a sample for imaging, basic bright field illumination is provided by the auxiliary flexi-lamp which may be tilted to give oblique illumination contrast to specimens, Figure 2A to A′′. The auxiliary flexi-lamp is also useful when assembling and aligning the instrument. For imaging proper, multiple transmission illumination modes are achieved using an 8×8 white LED array placed in the pupil plane (Liu *et al*., 2014). Using this condenser-free approach, it is possible to achieve modes analogous to traditional, condenser-based illumination (Webb, 2015). Illuminating all the LEDs provides basic bright field illumination (Figure 2B). Oblique illumination can be achieved by selectively illuminating strips of pixels from the different sides of the array (Figure 2B′). More advanced techniques are also possible, such as pseudo-phase contrast which is achieved by taking two oblique images and calculating the normalised difference between them (Figure 2B′′). Such post-acquisition contrast enhancement can be achieved automatically by the on-board processing of the microscope software. Another advanced technique, dark field illumination, is achieved by illuminating LEDs which fall outside of the numerical aperture (NA) of the chosen objective (Figure 2B′′′).

**Figure 2:**
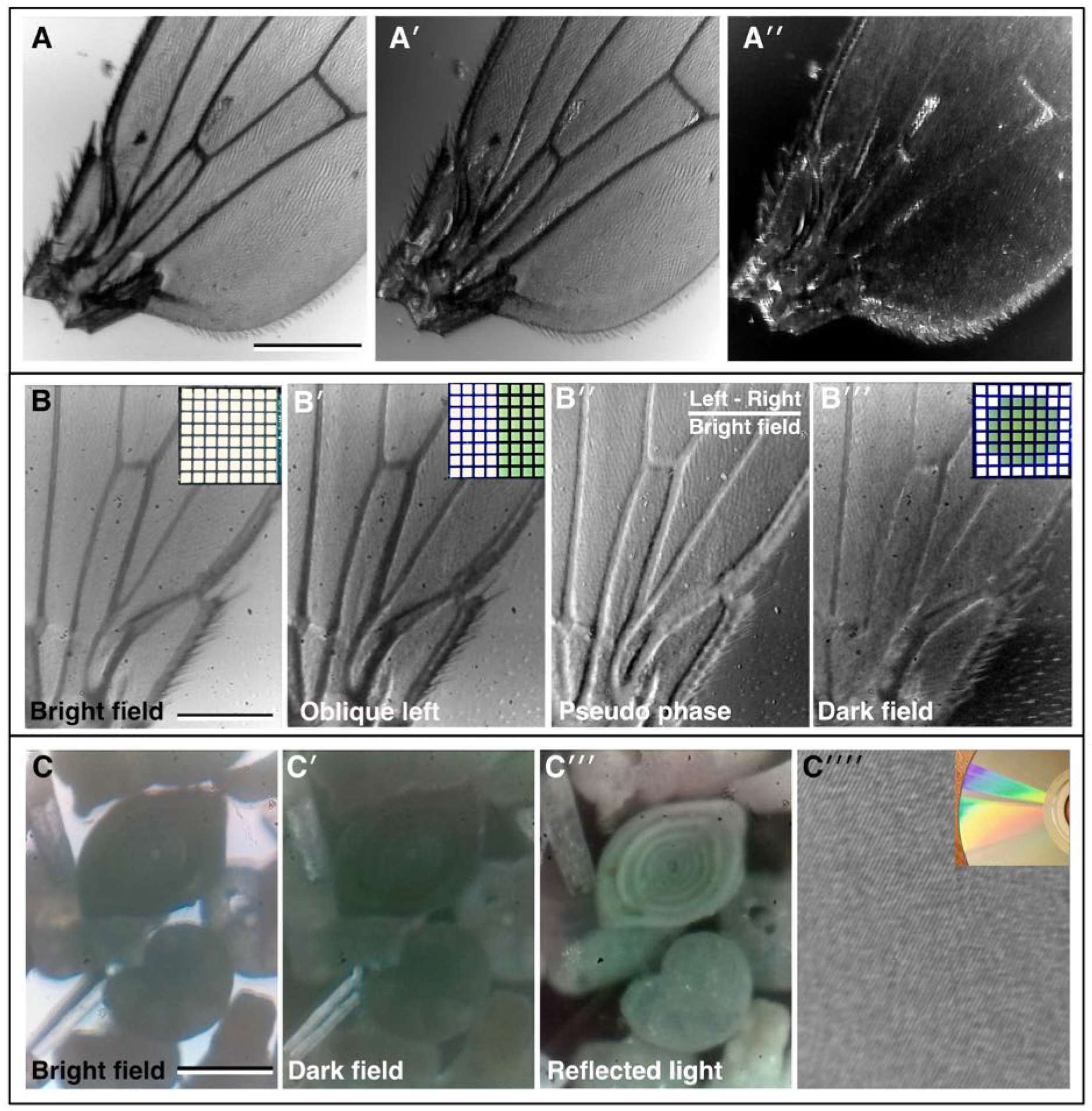
Bright field imaging modes. (**A-A′′**) A fruit fly (*Drosophila melanogaster)* wing imaged under direct illumination bright field (**A**) and different angles of oblique illumination (**A′, A′′**) using the auxiliary flexi-lamp illumination. (**B-B′′**) A fly wing imaged under different contrast methods using the bright field LED array: (**B**) bright field, (**B′**) oblique illumination, (**B′′**) dark field, (**B′′′**) pseudo-phase. Inset in B-B **′′′**, the LED configuration used to generate the different contrast effects in each case. (**C-C′′′**) Imaging a “fossil sand” sample from Dog’s Bay (Donegal, Ireland) in bright field; (**C**) dark field and (**C′′**) by reflected light. Several different microfossils are clearly visible. (**C′′′′**) Reflected light imaging of a completely opaque sample: the tracks on a DVD, inset, photograph of the DVD showing the reflection grating colour pattern caused by the tracks. (Scale bars = 0.3 mm in A, 0.22 mm in B and 0.25 mm in C.)

Microscopi, therefore, although superficially very different to a conventional teaching microscope, is still highly suitable for teaching the basics of optics, such as resolution, contrast and the concept of conjugate planes. The optical path can be customised to allow additional imaging modes such as polarised light imaging, by introducing polarising filters before and after the specimen, and reflected light imaging, using a ring of RGB LEDs beneath the objective (comparison in Figure 2C-C′′′ and C′′′′). The LED ring additionally allows the possibility of multiple Rheinberg illumination patterns (Rheinberg, 1896), which may be implemented by illuminating sectors of the LED ring with different colours.

Bright field imaging modes can reveal a wealth of information about a sample, although identification of specific features may be limited. The use of selective stains and colour imaging can provide far more specific information. Figure 3A shows an image of a typical histology slide of fixed tissue, in this case mouse aorta stained with Oil-Red-O which labels lipids, while Figure 3B-B′ shows a stained section of plant root. More recently, specific labelling of specimens has been revolutionised by fluorescence imaging of fluorescent-dye tagged probes. In fluorescence, molecules emit light at a specific visible wavelength (colour) upon illumination with light of a shorter wavelength. Using these fluorescent probes, specific features of the sample are highlighted by the bright fluorescence emission against a dark background. Figure 3B and B′ are images of dried-down spots of bright fluorescent green and red dyes respectively, both imaged with the same blue LED excitation and a long pass emission filter. Figure 3C shows the weaker native autofluorescence of pollen grains. To set up and align the fluorescence illumination, a test slide of fluorescent highlighter stained lens tissue was used (See Materials and Methods and Supplemental Figure S2).

**Figure 3:**
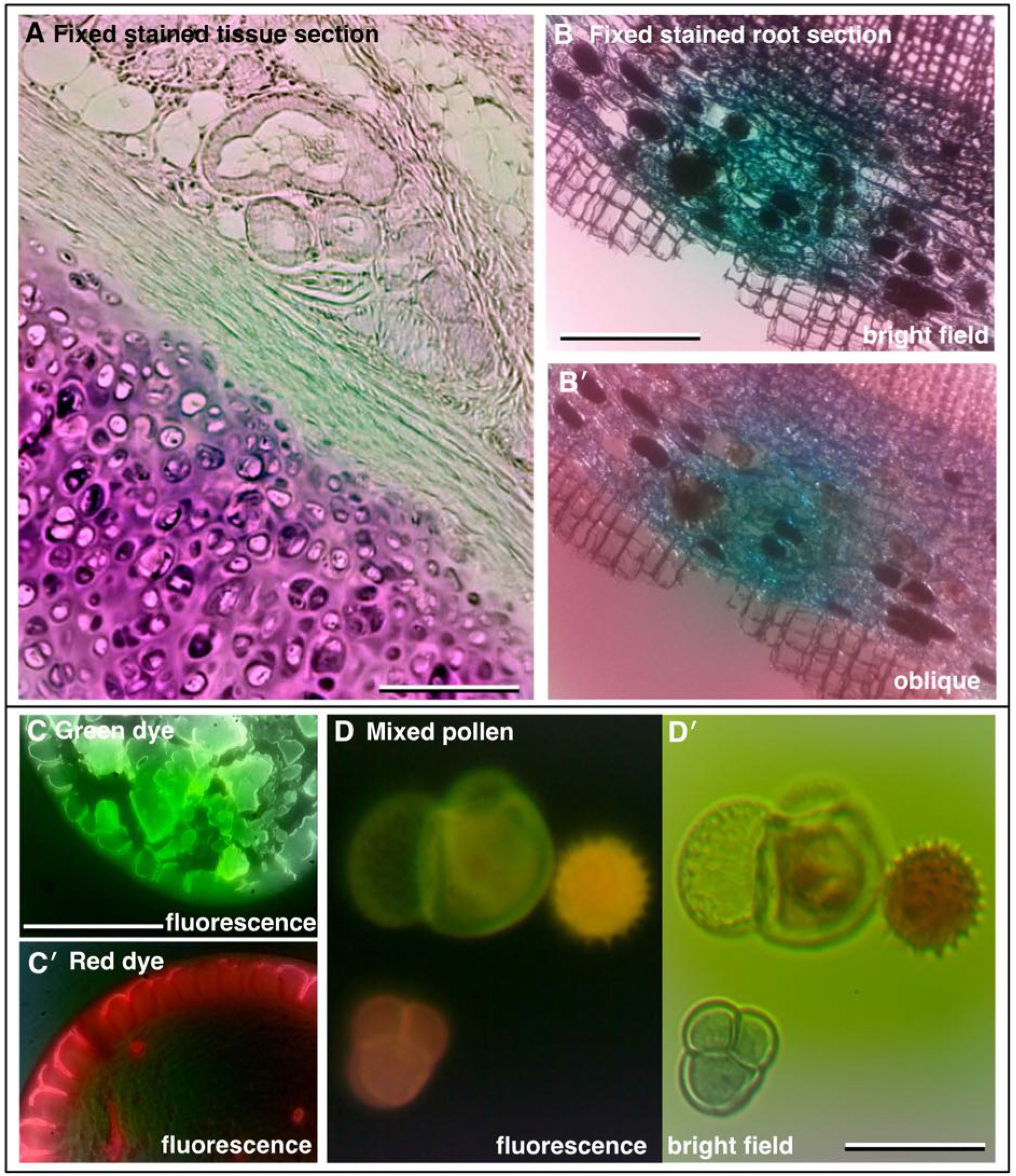
Increasing contrast with colour and fluorescence imaging. (**A**) Fixed and sectioned animal tissue, stained with Oil-Red-O showing a range of structures. (**B**) Fixed, sectioned and stained root of a plant (Ginko) showing the different cell layers. (**C, C′**) Fluorescence images of adjacent dried down puddles of green and red dye, respectively. (**D**) Autofluorescence of pollen grains and (**D′**) corresponding colour bright field image. (Scale bars = 0.20 mm in A, 0.35 mm in B, 0.35 mm in C and 0.04 mm in D.)

### Easy Navigation of Samples with Motorised Focus and XY Flexure Stage

One of the big challenges in using a microscope is finding and navigating the sample. For a teaching and outreach instrument it is critical that this initial step is made quick and easy. A major departure of Microscopi from traditional microscopes is the lack of eyepieces to assist in locating and orienting the sample. This allows for a simpler more compact overall design. To overcome the potential difficulty for users due to the lack of eyepieces, samples are navigated through the responsive motor-driven focus and XY stage monitored via a live feed from the system’s digital camera.

The motorised focus (Z-drive) of Microscopi is based on an M4 screw directly driving a 3D printed rail/slider to raise and lower the imaging lens. This approach provides a relatively large focusing range with well-defined increments provided by a stepper motor and little XY drift during focus. We were easily able to record a relatively large 3D sample such as a fly head (*Drosophila melanogaster*) as shown in Figure 4 and Supplemental Movies 1A, B. Automated focus control offers further options including reconstructing an extended focus or 3D image view of a specimen and automated autofocus functionality.

**Figure 4:**
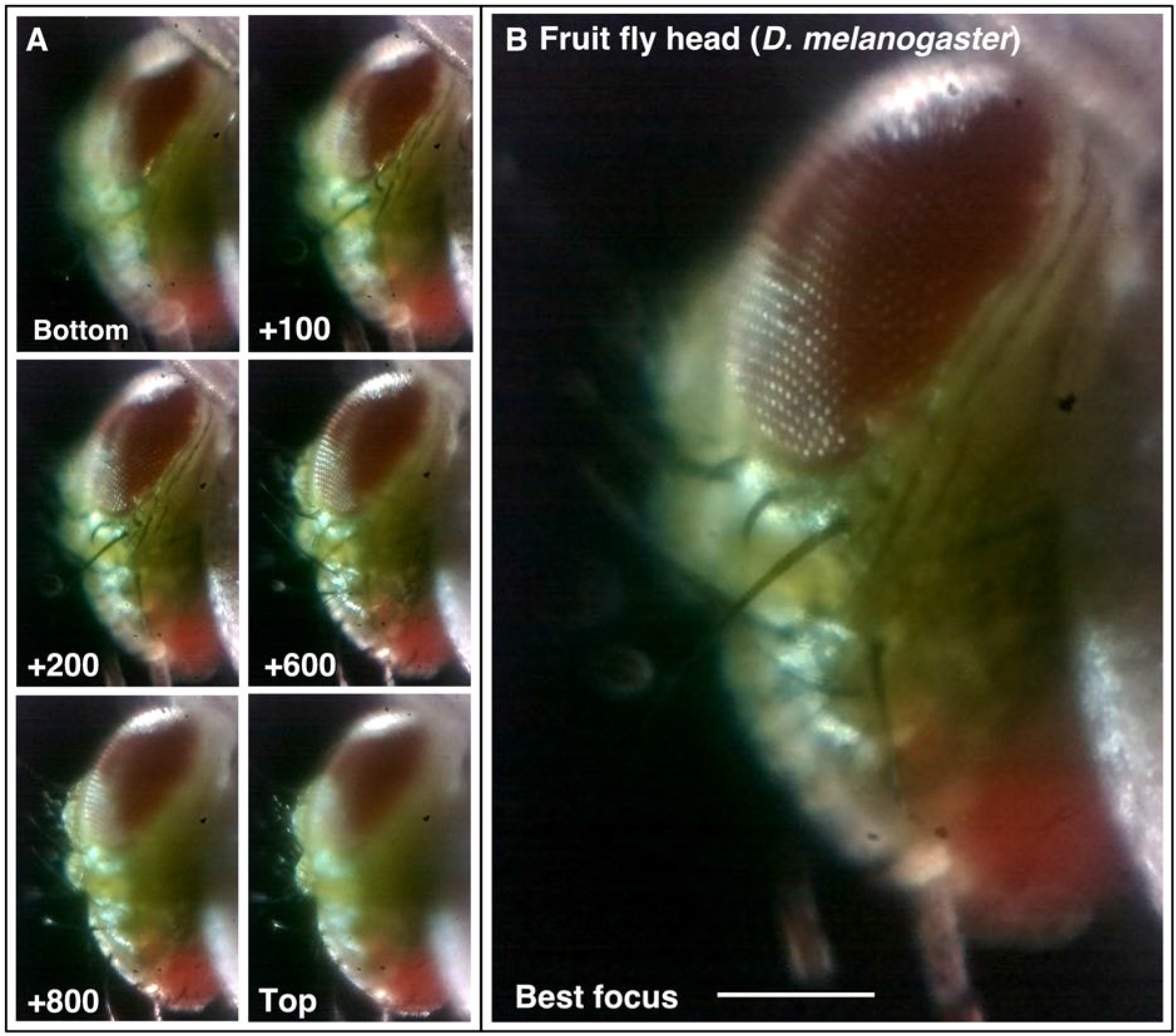
Motorised focus. (**A** and **B**) Using the motorised Z-drive to collect a focal series of a fly head (*Drosophila melanogaster*) to find the point of best focus. Z-units are arbitrary motor units, approximately 100 units/0.1 mm. See Supplementary Movie-1. (Scale bars = 0.15 mm in B.)

Navigation of the sample in the horizontal plane (XY directions) is achieved by a simple, elegant, 3D printed flexure-stage which mimics real-world precision stages. The sample platform is held by four flexures, each incorporating a push plate. The stage movement is driven from motors with an integrated captive nut. A drive-rod protrudes from the motor and pushes directly against the push plate to displace the flexure. The flexure design provides backlash-free motion with little Z-drift, although the actual trajectory of travel is over a slight arc due to the constraint of the bent flexures. Symmetric flexures and parallelogram shape help to alleviate this (Bankar, 2017). With the motorised flexure-stage it is possible to both easily navigate across larger samples and generate a single composite image as shown in the tiled 5×5 panel view of the sectioned stained animal tissue histology slide in Figure 5. Fine control of XY and Z also make it possible to use higher powered objectives facilitating imaging at different scales (Supplemental Figure S3).

**Figure 5:**
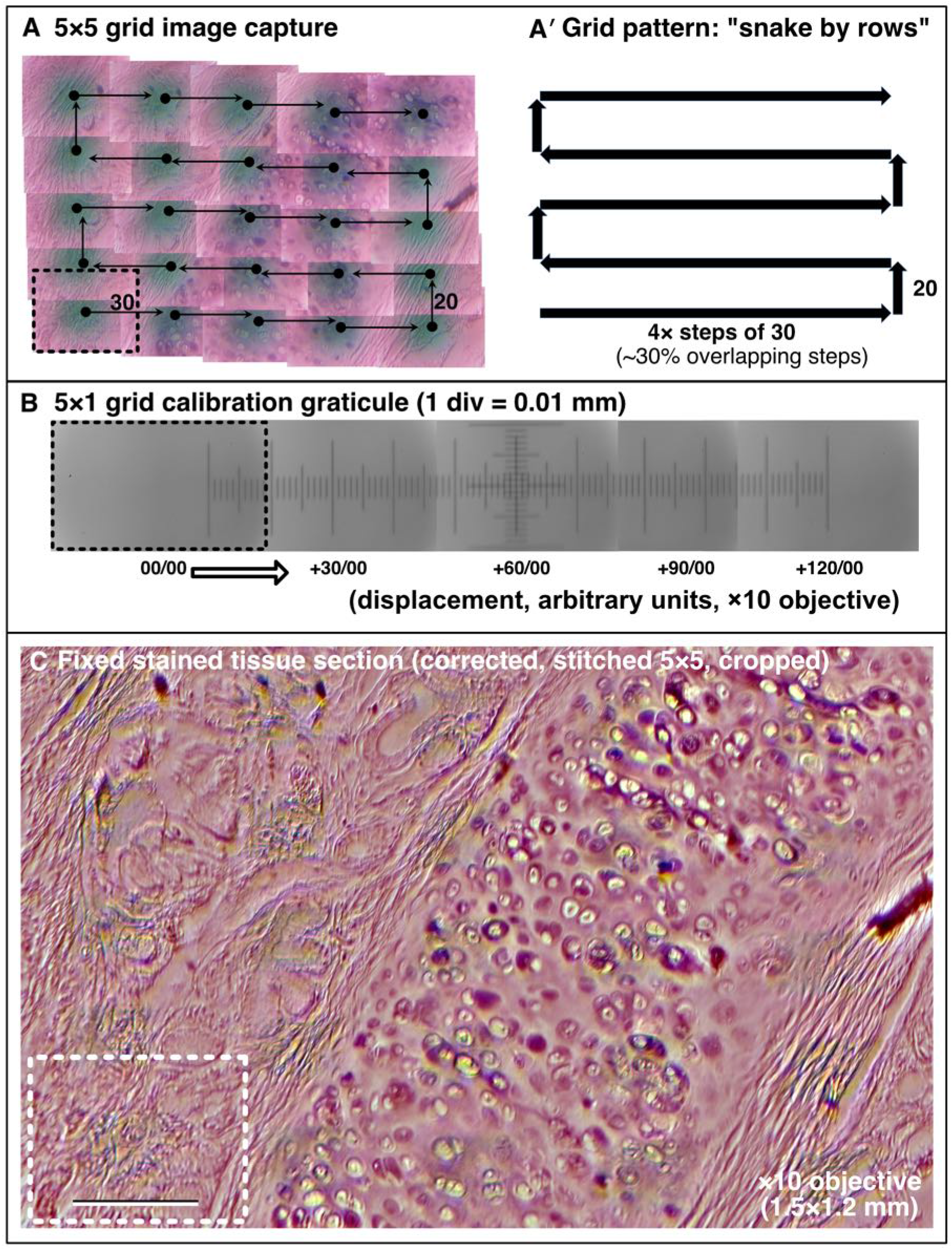
Motorised XY sample navigation. (**A** - **C**) Using the motorised flexure-stage and an automated stitching algorithm to extend the field of view to cover a large sample. (**A, A′**) Example of a 5×5 image collection pattern of stage movements following a “snake by rows” grid with 30% tile overlap. (**B**) Graticule scale imaged with the same ×10 objective indicating the dimensions of movement. (**C**) Final processed stitched image of a fixed sectioned animal tissue histology slide stained with Oil-Red-O covering a 1.5×1.2 mm field. Dashed box = 1 image tile. (Scale bar in C = 0.2 mm.)

### An Exercise in Control – the GUI and Underlying Code

Microscopi is a motorised microscope with a digital camera. These features lend themselves to automation, computer control and image processing. We have taken advantage of this to make Microscopi an integrated imaging system such that a user can navigate or scan a sample to produce extended focus images, tiled composite views (Figure 5C) and time-lapse movies (Supplemental Movies 2A, B) remotely without having to open up the microscope and adjust hardware directly.

Microscopi is controlled by dedicated Python-based control software operating on an on-board Raspberry Pi computer (See Materials and Methods). We take advantage of the prevalence of electronic devices such as smartphones, desktop computers and laptops to run the Graphical User Interface (GUI) to allow a user to interact with the microscope (Figure 6). The onboard Raspberry Pi acts as a Wi-Fi hotspot to which devices connect. Once a device, such as a smartphone, has connected to the local Wi-Fi any web page address of the form “http://anything” redirects to the Microscopi GUI web interface. This enables full control of the microscope automation, such as sample navigation, focus, and illumination. In addition, images can be captured, viewed, and downloaded.

**Figure 6:**
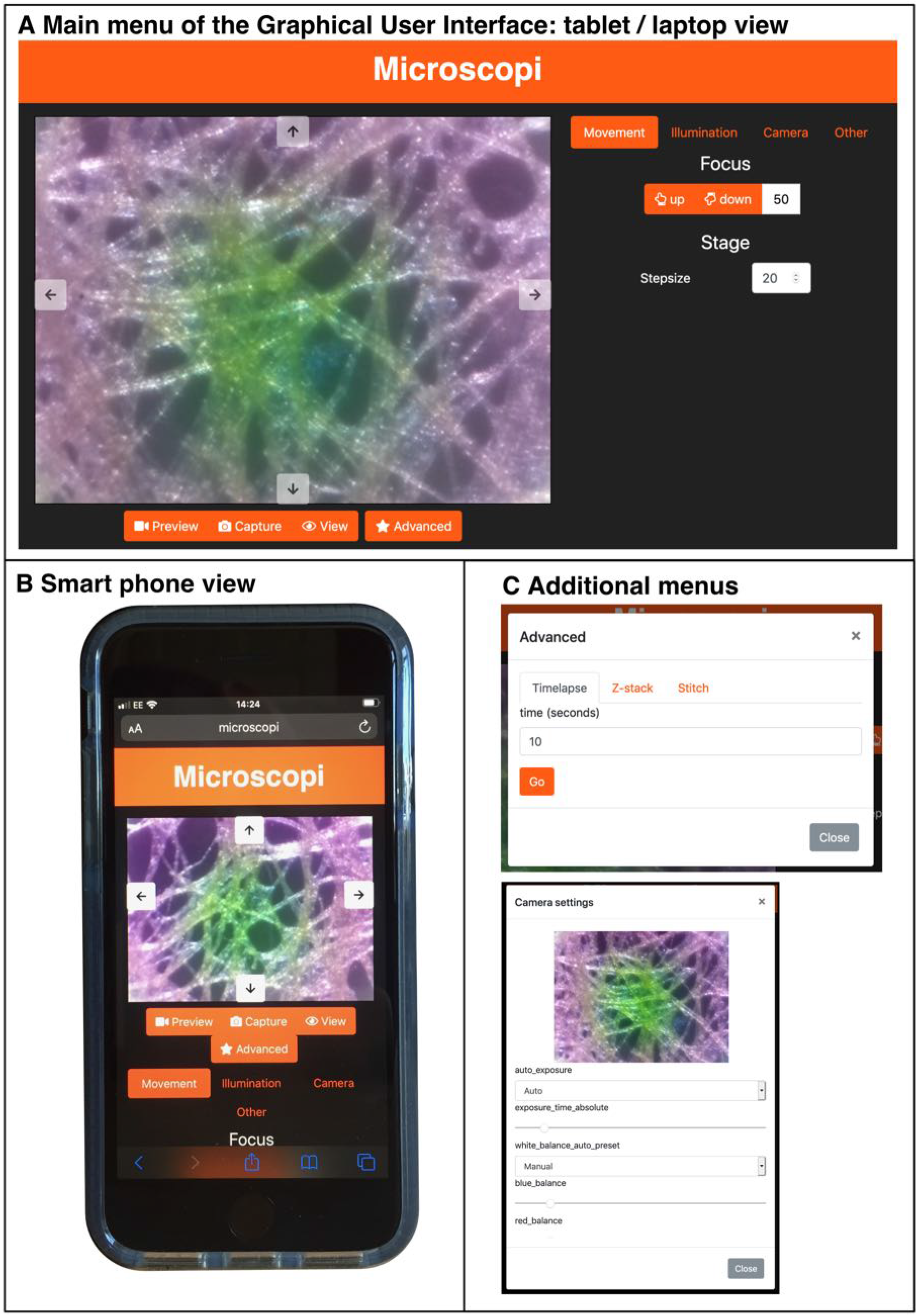
The Microscopi Graphical User Interface. (**A**) Layout of the Microscopi graphical user interface using a desktop/laptop browser. (**B**) Layout of the graphical user interface as used on a smartphone. (**C**) Example of additional modal popups.

It is possible to achieve a higher level of automation with relatively simple Python coding to provide additional functions which can be configured and launched from the GUI. The “automated Z-series” collects images though a sample with a user defined step size and range. We have also implemented a simple autofocus mechanism. The most basic autofocus uses a microswitch to define a fixed home position. From the home position an expected sample focus is reached by a fixed offset for each lens. Fine-tuning of this focus may be achieved by subsequently imaging through the sample in small Z steps, assessing the image series against a sharpness or focus metric and navigating to the Z-position with the highest score. Automation can also be applied to increase the effective field of view across XY by incrementing the XY stage movement in a “snake by rows” pattern (Preibisch *et al*., 2009) collecting a 3×3 or a 5×5 image tile array, followed by image-content based stitching. GUI based control of the LED array illumination pattern allows a user to explore different contrast modes to view a sample, and combining any of these protocols further increases the scope for automation.

While operating the microscope from a smartphone is preferable in many usage scenarios it is less ideal for customising the code to add additional features. Microscopi can be controlled by a keyboard/mouse/monitor plugged into the accessible ports of the Raspberry Pi allowing access to the code running directly on the instrument. The simple Python modules that permit extensive microscope control and experimentation from the GUI can be called or customised and added to. These new features can be subsequently incorporated into the GUI. The ease with which both the hardware and software of Microscopi can be modified and customised further adds to its value in teaching at various education levels and in customisation to niche applications decided by the user.

## PERSPECTIVE

### A Versatile Imaging System for Teaching, Outreach and Fieldwork

Our goal in developing Microscopi has been to provide a widely accessible, fully functional automated imaging system, supporting a wide range of interdisciplinary science teaching activities in schools and colleges. A key future objective is to develop a community of users by freely distributing detailed assembly instructions, control software and worked examples of practical exercises. There is also a need for accessible, easy-to-use and inexpensive fluorescence imaging technology, not only for teaching but in real-world applications for medical or veterinary diagnosis or assessment of water quality. Our future goals are to adapt the Microscopi design to address these needs and establish funding to support the generation and distribution of hardware in kit form.

## MATERIALS AND METHODS

### Design Procedures and 3D Printing

The layout of Microscopi and 3D components were designed in the freely available CAD package OpenSCAD and exported as STL files. Components were printed on a Lulzbot Taz 5 or Ultimaker S5 printer. Printing parameters were varied according to the requirements of the component, for example, the moving parts of the focus mechanism were printed on finer settings. All printed components were made from PLA filament which offered the best compromise of strength, reproducibility and ease of use. Used PLA is unfortunately not currently widely recycled or widely compostable so disposing of misprints or test prints and the disposal of the microscope when it reaches end-of-life should be carefully considered, or alternative materials tested for suitability. All design files and exported STL files may be found at https://doi.org/10.5281/zenodo.3701602 as well as part numbers, detailed build instructions and updates. Design drawings for incorporation into figures were produced in the free Blender 2.82 drawing and design software.

### Assembly of Components and Acceptance Testing

The build procedure is documented in the Supplementary Methods “Assembly Guide” hosted online at https://doi.org/10.5281/zenodo.3701602. The Supplementary Methods also includes operating instructions and a series of acceptance tests to assist in commissioning the build, as well as a brief trouble-shooting guide and safety documentation. We strongly advise adult supervision of anyone under the age of 18 during the build process and for operation in fluorescence imaging mode. Following the instructions, Microscopi is relatively simple to build from its component parts, most of which require only nuts and bolts to assemble. Hot glue is recommended for adhering parts together as it is non-toxic and easily reversible. Assembling the electronics and wiring is more technically challenging than the other tasks and requires access to more specialised tools (for example: wire snips, wire stripper and soldering iron). As far as possible soldering has been replaced with “solder free’ options. General instructions on how to assemble cables and use the various tools may be readily found online.

### Safety Considerations

Microscopi is intended to be an easy-to-use, accessible instrument for use outside the science lab environment and by the public. However, the optional LED used for fluorescence illumination is bright, so staring into the beam should be avoided. To prevent accidental exposure, the lid should be fitted with an interlock microswitch which disables the fluorescence LED when opened. We also recommend that the outside casing of the instrument is clearly marked with the warning stickers supplied with the fluorescence LED. A warning label also features in the Components and Assembly Guide. Microscopi is modular, therefore, for younger user groups, we recommend only installing the bright field 8×8 white LED, LED ring-light and the flexi-lamp.

### Software Development and Implementation

Microscopi is driven by several interacting software components running under Raspbian, a Debian-based Linux operating system. In particular, the majority of the microscope control code and GUI are implemented in a custom Python package. The Python package broadly serves two functions. The first is control of the hardware components of the microscope, such as the motors, illumination and camera. In this capacity, the python package can also be used independently from the GUI to write simple automated procedures for the microscope, such as Z-stacks. The second function is to implement the GUI. GUI implementation is achieved by running a webserver (based on Flask) which serves a webpage, containing the GUI, to clients. Running both the control and web server using Python allows simple interaction between the two. The basic Raspbian operating system is configured with additional software and service files to set up the wireless hotspot and to redirect connected clients to the Microscopi GUI served by the Python package. Detailed instructions on how to download, install and implement the Microscopi software are provided in the documentation https://doi.org/10.5281/zenodo.3701602 and the code is available at https://github.com/micronoxford/microscopi.

### Sample Preparation

A variety of samples were sourced for testing, including commonly found materials and pre-prepared commercially available slide sets (see Table of Resources). Samples, such as pond water and beach sand, were collected and viewed in plain 35 mm plastic sample dishes or 35 mm plastic dishes with a glass coverslip bottom. Other samples, such as moss leaves and other plant fragments, were mounted in a drop of rainwater on a glass slide beneath a No. 1.7 22×22 mm coverslip. To avoid crushing delicate samples, small sections of double-sided sticky tape were used between the slide and coverslip. Fly parts were obtained from individual flies from lab-cultured *Drosophila*, killed by extended exposure to CO_2_. Mineral slides were kindly provided by Owen Green (University of Oxford, Oxford Earth Sciences). Oil-Red-O stained mammalian tissue section slides were kindly provided by Gillian Douglas (University of Oxford WTCHG).

A standard fluorescent test slide was produced by colouring an approximately 10 mm square section torn from a sheet of lens cleaning tissue, (alternatively a single ply of toilet paper could be used), mounting dry on a slide beneath a 22 mm square coverslip and sealing with nail varnish, as described in the Supplementary Methods (hosted online at https://doi.org/10.5281/zenodo.3701602).

### Image Collection and Processing

Images were collected on Microscopi using either ×3, ×4, ×10, or ×40 lenses. For some automated features, on-board post acquisition image processing is performed using custom Python scripts. Any post acquisition image manipulation or processing applied for the preparation of images for figures was carried out in the freely available Fiji V2.0.0 image analysis software (for example, see Supplementary Figure S4). Image data was archived in OMERO V5.3.5 (Open Microscopy Environment).

### Data and Software Availability

Software: https://github.com/micronoxford/microscopi.

Microscopi Assemby Manual: https://doi.org/10.5281/zenodo.3701602 (Updates will be made available through the Zenodo site)

## AUTHOR CONTRIBUTIONS

MW: Software, Methodology, Visualization, Validation, Conceptualization, Writing (original draft), Writing (review and editing). AJ: Methodology, Validation, Writing (review and editing). IMD: Conceptualization, Writing (review and editing). MJB: Conceptualization, Project administration, Funding acquisition, Writing (review and editing). ID: Conceptualization, Funding acquisition, Project administration, Writing (review and editing). RMP: Methodology, Investigation, Visualization, Validation, Conceptualization, Project administration, Writing (original draft), Writing (review and editing).

## ACKNOWLEDGEMENTS

Special thanks to Catrina F. Parton for testing the system, providing user feedback and collecting image data. We would also like to thank Tanja Kollakowski, Mick Phillips, Nicholas Hall and Darragh Ennis for helpful discussions, Ee Zhuan Chong and Jun Guan for assistance with methodology and Martin Hailstone for advice on image analysis. This work was supported by: Wellcome Strategic Awards 091911/B/10/Z and 107457/Z/15/Z) to ID, a Wellcome Trust Senior Research Fellowship (081858) to ID; a Wellcome Investigator Award (209412/Z/17/Z) to ID; a John Fell Fund Award 141/020 to ID ; an EPSRC IAA grant EP/K503769/1 to MJB.

## TABLE OF RESOURCES^*^

TABLE OF RESOURCES\*
| REAGENT or RESOURCE | SOURCE/MANUFACTURER | IDENTIFIER |
| --- | --- | --- |
| **Materials** |  |  |
| General hardware components & Tools | Farnell/CPC/OneCall<br>RS |  |
| General optics components | Thorlabs |  |
| lenses (160, DIN) | OPTIKA, MOTIC | https://www.raig.co.uk M164, M165, M166, M167 |
| Electronic components | Farnell/CPC/OneCall<br>RS<br>Hobbytronics |  |
| LED arrays | PiHut<br>Pimoroni | ADA1857, ADA2856 |
| LED adjustable spotlight | Thorlabs | PSX501 |
| **3D Printing** |  |  |
| Lulzbot Taz 5 | Aleph objects |  |
| Ultiimaker S5 | Ultimaker | https://ultimaker.com/3d-printers/ultimaker-s5 |
| Ultimaker Cura 4.5 | Ultimaker | https://ultimaker.com/software/ultimaker-cura |
| Ultimaker PLA filament | RS | 134-8185 |
| **Software Packages** |  |  |
| Python | www.python.org |  |
| Raspbian | https://www.raspbian.org/ |  |
| Fiji, ImageJ (V2.2.0) | https://fiji.sc<br>http://imagej.nih.gov/ij | Schindelin *et al.*, (2012) |
| OMERO V5.3.5 | openmicroscopy.org/omero/ | Allen *et al.*, (2012) |
| OpenSCAD | https://www.openscad.org |  |
| Blender | https://www.blender.org |  |
| Cura | https://ultimaker.com/software/ultimaker-cura |  |
| **Samples** |  |  |
| Prepared slides | https://www.ukge.com | Prepared slides sets |
| Slides and coverslips | https://www.ukge.com | MC0300, MC0350 |
| Fluorescent dye Fluorescein | eBay | Fluorescein LTC - Acid Yellow 73 |
| Fluorescent dye Rhodamine | eBay | Rhodamine B |
| Fluorescent highlighter Pens | Office Depot (Foray) | eBay |
| Lens cleaning tissue (for cameras) | eBay |  |
| Clear Nail Varnish | eBay | Avon Mark |
| Nice-day Double sided tape | https://www.viking-direct.co.uk | ND180492 |
| 35mmx10mm Plastic Petri dishes + lid | eBay |  |

\* A full listing of materials for assembling the instrument may be found in the Supplementary Methods online at https://doi.org/10.5281/zenodo.3701602.

## SUPPLEMENTARY MATERIAL

Supplementary Methods: hosted online at https://doi.org/10.5281/zenodo.3701602

Supplementary Figure 1: Microscopi imaging system

Supplementary Figure 2: Lens tissue test sample

Supplementary Figure 3: Imaging at different scales

Supplementary Figure 4: Image colour artefact correction

Supplementary Movie-1A: Focal series of a fly head

Supplementary Movie-1B: XY Aligned focal series of a fly head

Supplementary Movie-2A: Timelapse of *Daphnia* (waterflea), 30s

Supplementary Movie-2B: Timelapse of onion epidermis cytoplasmic streaming, 30s

**Figure S1:**
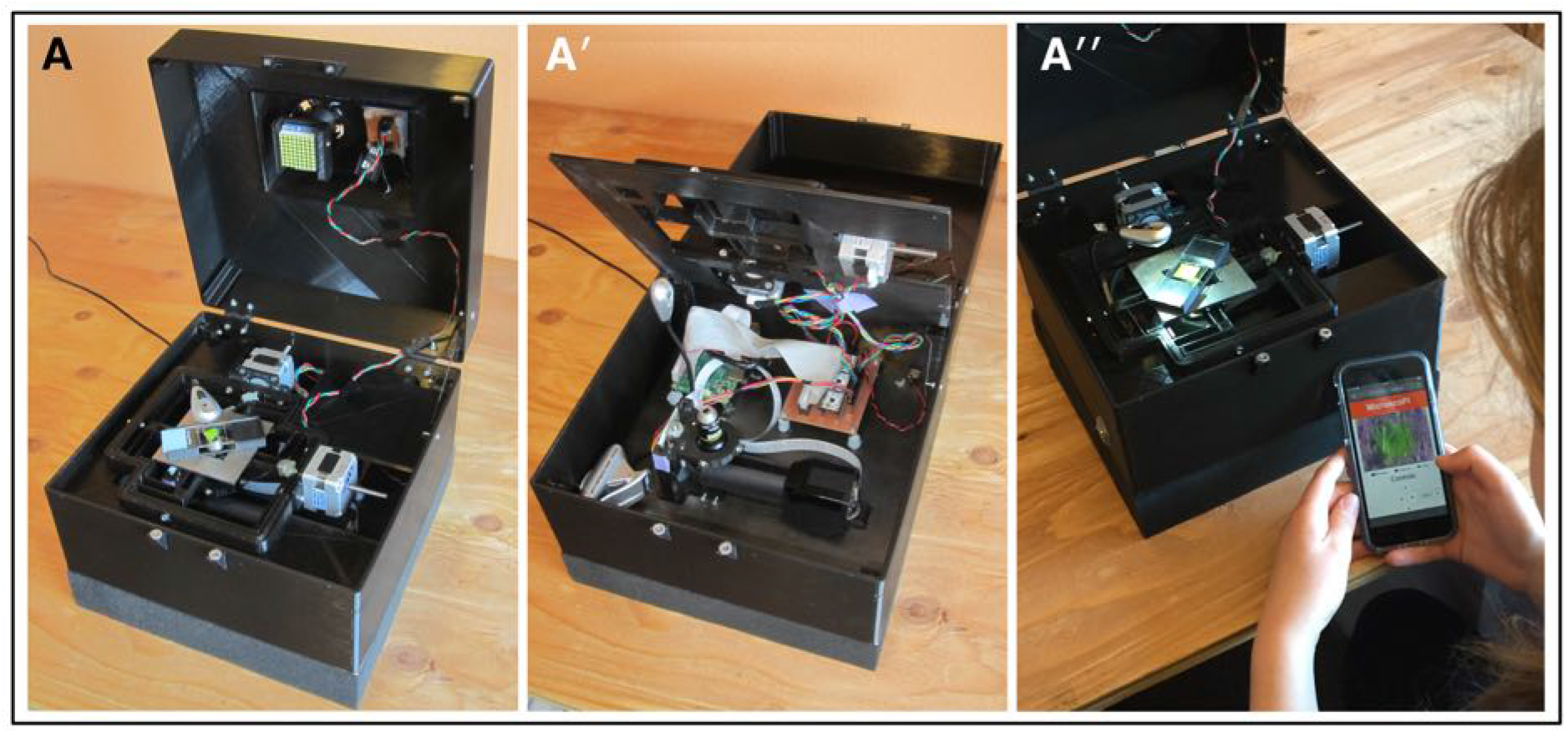
The Microscopi imaging system. (**A-A′′**) The fully assembled unit (**A**); with the hinged sample platform open showing the easy access to the light path and electronics (**A′**); in use from a smartphone (**A′′**).

**Figure S2:**
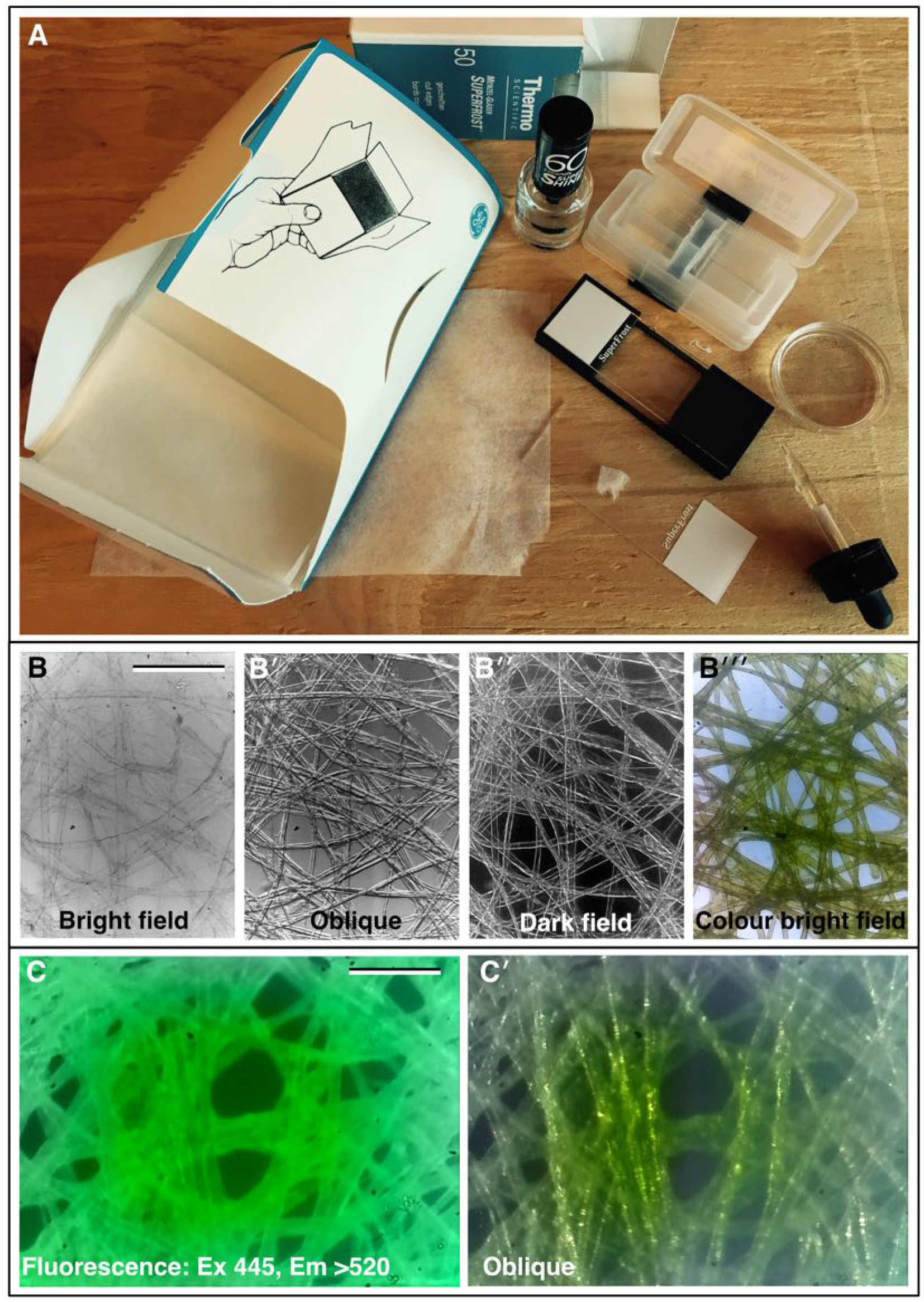
Lens tissue test sample. (**A**) Layout of the materials required to produce a simple lens tissue test sample, refer to the Supplementary Methods hosted online at https://micronoxford.com/microscopi for full instructions. (**B**-**B′′′**) Examples images of lens-tissue test slides taken on Microscopi using the bright field LED array and ×4 lens NA 0.1: (**B**) basic bright field, (**B′**) oblique illumination, (**B′′**) dark field, (**B′′′**) colour bright field of a fluorescent highlighter pen coloured section of lens tissue mounted dry. (**C**-**C′**) Fluorescence imaging of a highlighter pen coloured section of lens tissue mounted dry and corresponding oblique illumination bright field, ×4 lens NA 0.1. (Scale bars = 0.25 mm in B; 0.15 mm in C.)

**Figure S3:**
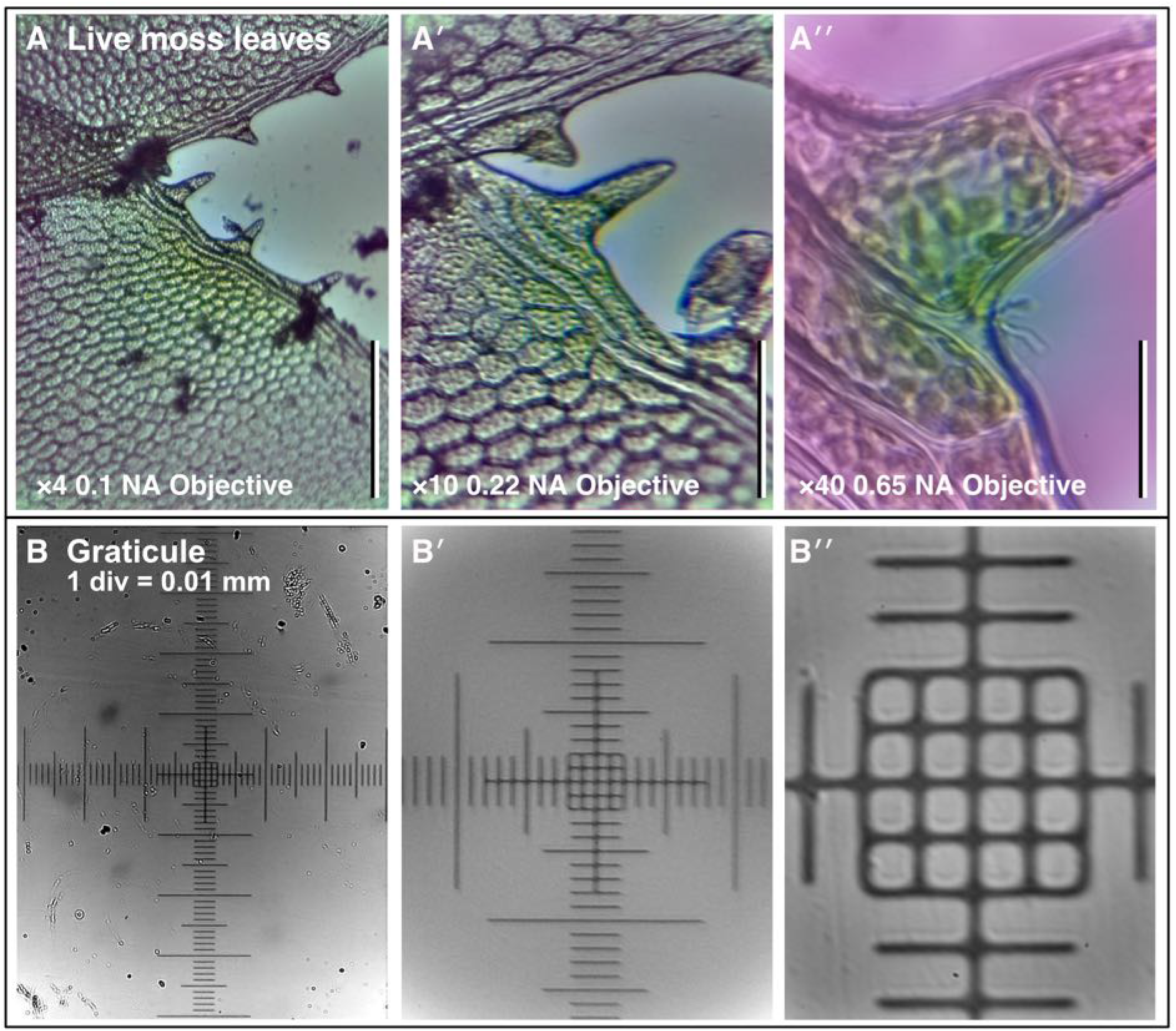
Imaging at different scales. (**A-A′′**) Live moss leaf sample imaged with ×4, ×10 and ×40 objectives, respectively. (**B-B′′**) Corresponding calibration images of a graticule slide 1 div = 0.01 mm for the three different lenses. (Scale bars = 0.3 mm in A, 0.2 mm in A**′** and 0.03 mm in A**′′**.)

**Figure S4:**
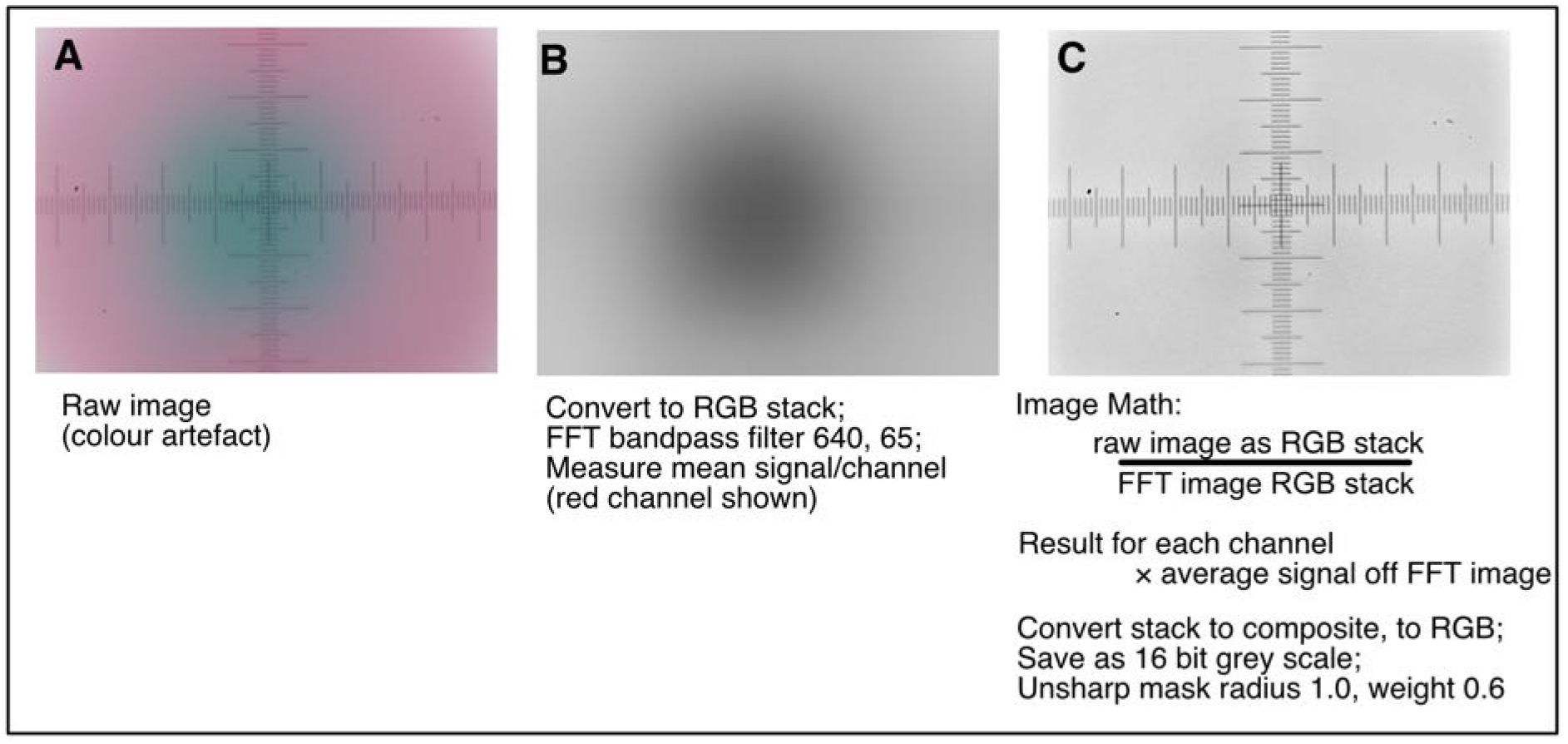
Image colour artefact correction in Fiji. (**A-A′′**) Post acquisition image processing in Fiji (ImageJ) to correct a colour artefact commonly found in the Pi camera V2 used without the Pi camera lens (https://www.arducam.com/docs/cameras-for-raspberry-pi/native-raspberry-pi-cameras/lens-shading-calibration/). (\***Note**: updates to the camera software are currently underway to address this issue https://libcamera.org/.)

## REFERENCES

Allan, C, Burel, J-M., Moore, J, Blackburn, C, Linkert, M, Loynton, S, Macdonald, D, Moore, WJ, Neves, C Patterson, et al. (2012). OMERO: flexible, model-driven data management for experimental biology. Nat Methods 9, 245–253. https://doi.org/10.1038/nmeth.1896

Bankar, SG (2017). Modeling and Design Analysis of XY Flexure Stage Mechanism. International Journal of Engineering Research & Technology 6 (4), 899–904. (IJERTV6IS040686)

Bogoch, II, Andrews, JR, Speich, B, Utzinger, J, Ame, SM, Ali, S. & Keiser J (2013). Mobile Phone Microscopy for the Diagnosis of Soil-Transmitted Helminth Infections: A Proof-of-Concept Study. The American Journal of Tropical Medicine and Hygiene 88 (4), 626 –629. https://doi.org/10.4269/ajtmh.12-0742

Breslauer, DN, Maamari, RN, Switz, NA, Lam, W. & Fletcher, DA (2009). Mobile Phone Based Clinical Microscopy for Global Health Applications. PLoS ONE 4(7): e6320. https://doi.org/10.1371/journal.pone.0006320

Collins, J, Knapper, J, Stirling, J, Mduda, J, Mkindi, C, Mayagaya, V, Mwakajinga, G, Nyakyi, P, Sanga, V, Carbery, D, White, L, Dale, S Jieh Lim, Z. Baumberg, J, Cicuta, P, McDermott, S, Vodenicharski, B and Bowman, R (2020). Robotic microscopy for everyone: the OpenFlexure microscope, Biomed. Opt. Express 11, 2447–2460. https://doi.org/10.1364/BOE.385729

Cybulski, JS, Clements, J, & Prakash M (2014) Foldscope: Origami-Based Paper Microscope. PLOS ONE 9(6): e98781. https://doi.org/10.1371/journal.pone.0098781

Diederich, B, Lachmann, R, Carlstedt, S, Marsikova, B, Wang, H, Uwurukundo, X, Mosig, A & Heintzmann, R (2020). UC2 – A Versatile and Customizable low-cost 3D-printed Optical Open-Standard for Microscopic imaging. bioRxiv 2020.03.02.973073; doi:https://doi.org/10.1101/2020.03.02.973073

Hergemoller, T & Laumann, D (2017). Smartphone Magnification Attachment: Microscope or Magnifying Glass The Physics Teacher 55, 361; https://doi.org/10.1119/1.4999732

Liu, Z, Tian, L, Liu, S, Waller, L (2014). Real-time brightfield, darkfield, and phase contrast imaging in a light-emitting diode array microscope. J Biomed Opt 19, 106002. https://doi.org/10.1117/1.JBO.19.10.106002

Orth, A, Wilson, ER, Thompson, J. & Gibson, BC (2018). Dual-mode mobile phone microscope using the onboard camera flash and ambient light. Scientific Reports, Volume 8, Article number: 3298. https://doi.org/10.1038/s41598-018-21543-2

Pirnstill, C. & Coté, GL (2015). Malaria Diagnosis Using a Mobile Phone Polarized Microscope. Scientific Reports volume 5, Article number: 13368. https://doi.org/10.1038/srep13368

Preibisch, S, Saalfeld, S & Tomancak, P (2009). Globally optimal stitching of tiled 3D microscopic image acquisitions. Bioinformatics (Oxford, England), 25(11), 1463–1465. https://doi.org/10.1093/bioinformatics/btp184

Rheinberg, J. (1896) On an addition to the methods of microscopical research, by a new way optically producing colour-contrast between an object and its background, or between definite parts of the object itself., J. R. Microsc. Soc.16, 373–388

Schindelin, J, Arganda-Carreras, I, Frise, E et al. (2012). Fiji: an open-source platform for biological-image analysis. Nat Methods 9, 676–682. https://doi.org/10.1038/nmeth.2019

Simon, J. & Comastri, S (2015). “The compound microscope: optical tube length or parfocalization?”. European Journal of Physics, vol. 26. https://doi.org/10.1088/0143-0807/26/6/018

Switz NA, D’Ambrosio MV & Fletcher DA (2014). Low-Cost Mobile Phone Microscopy with a Reversed Mobile Phone Camera Lens. PLOS ONE 9(5): e95330. https://doi.org/10.1371/journal.pone.0095330

Tristan-Landin SB, Gonzalez-Suarez AM, Jimenez-Valdes RJ & Garcia-Cordero JL (2019) Facile assembly of an affordable miniature multicolour fluorescence microscope made of 3Dprinted parts enables detection of single cells. PLoS ONE 14(10): e0215114. https://doi.org/ 10.1371/journal.pone.0215114

Wei, Q, Qi, H, Luo, W, Tseng, D, Ki, SJ, Wan, Z, Göröcs, Z, Bentolila, LA, Wu, T-T, Sun, R and Ozcan, A (2013). Fluorescent Imaging of Single Nanoparticles and Viruses on a Smart Phone ACS Nano, Vol 7 (10), pp 9147–9155. https://doi.org/10.1021/nn4037706

Webb, KF (2015). Condenser-free contrast methods for transmitted-light microscopy. J Microsc 257, 8–22. https://doi.org/10.1111/jmi.12181

Zheng, G, Kolner, C, & Yang, C (2011). Microscopy Refocusing and Dark-Field Imaging by Using a Simple Led Array. Opt Lett 36, 3987–3989. https://doi.org/10.1364/OL.36.003987

Zhua, H, Sencana, I, Wonga, J, Dimitrova, S, Tsenga, D, Nagashimaa, K and Ozcan A (2013). Cost-effective and rapid blood analysis on a cell-phone. Lab Chip, 13, p1282. https://doi.org/10.1039/c3lc41408f

